# Extracellular vesicles carrying surface-anchored adiponectin prevent obesity-related metabolic complications by enhancing insulin sensitivity

**DOI:** 10.1101/2025.09.04.674183

**Authors:** Alexia Blandin, Josy Froger, Margot Voisin, Maëlle Lachat, Grégory Hilairet, Katarzyna Polak, Julie Magusto, Lisa Meslier, Mikaël Croyal, Mathilde Gourdel, Quentin Massiquot, Julien Chaigneau, Maxime Carpentier, Lionel Fizanne, Xavier Prieur, Cédric Le May, Jérôme Boursier, Bertrand Cariou, Robert Mamoun, Bernadette Trentin, Soazig Le Lay

**Affiliations:** Nantes Université, CHU Nantes, CNRS, INSERM, l’institut du thorax, F-44000 Nantes, France; Université Angers, SFR ICAT, 49000 Angers, France; Ciloa, 356 rue Maurice Béjart, 34080 Montpellier, France; Nantes Université, CHU Nantes, Inserm, CNRS, SFR Santé, Inserm UMS 016, CNRS UMS 3556, Nantes, France; HIFIH, CHU Angers, Université Angers, SFR ICAT, 49000 Angers, France

**Author notes:** These authors equally contributed to this work (Alexia Blandin, Josy Froger).

**Keywords:** Extracellular vesicles, exosomes, obesity, type 2 diabetes, MASLD, MASH, biotherapy, bioengineering, EV-based therapy

## Abstract

Adiponectin (Adpn) is a potent insulin-sensitizing adipokine with therapeutic promise for type 2 diabetes (T2D) and metabolic dysfunction-associated steatohepatitis (MASH). Its clinical use is limited by challenges in producing stable, bioactive high-molecular weight forms. Adipocyte-derived extracellular vesicles (EVs) naturally carry oligomeric Adpn on their surface, enhancing hormone stability and activity. Here, we engineered EVs displaying membrane-anchored Adpn (EV^PP-Adpn^) and control EVs lacking Adpn (EV^CTL^), and evaluated their metabolic effects in high fat diet (HFD)-induced obesity mice.

EV^PP-Adpn^ were purified from HEK293T cells stably transfected with a chimeric Adpn fused to a transmembrane domain and a pilot peptide (PP) directing it to EVs; EV^CTL^ were produced from non-transfected cells. HFD-fed male and female mice received intraperitoneal EV injections for six weeks.

EV^PP-Adpn^ improved glucose tolerance and insulin sensitivity, promoted adipocyte lipid storage through insulin-regulated lipogenesis and alleviated MASH features (liver steatosis, inflammation and fibrosis). EV^PP-Adpn^ lowered circulating ceramides and reduced FGF21, indicating improved hepatic metabolism, and activated AKT and AMPK pathways in liver and skeletal muscle, consistent with increased adiponectin signaling.

These results demonstrate that surface-anchored Adpn EVs restore tissue-specific insulin signaling and improve obesity-related metabolic dysfunctions, highlighting their potential as a novel biotherapeutic strategy for T2D and MASH.

## Introduction

The rising prevalence of type 2 diabetes (T2D) and its associated cardiometabolic complications underscores the need for innovative therapeutic strategies, especially those aimed at restoring insulin sensitivity. Despite major advances in glucose-lowering therapies, there remains a critical lack of disease-modifying approaches capable of simultaneously improving systemic insulin resistance and associated metabolic organ dysfunctions, including metabolic dysfunction-associated steatohepatitis (MASH).

Adiponectin (Adpn), an adipokine with insulin-sensitizing, anti-inflammatory, and cardioprotective properties is an attractive therapeutic target (1). High-molecular-weight (HMW) oligomers are recognized as the biologically active forms of the hormone (2; 3). Adpn oligomers signal through AdipoR1/2 to activate AMPK and PPARα-dependent metabolic pathways (4). AdipoR signaling also cross-talks with the insulin receptor and its downstream PI3K/AKT signaling pathways, thereby enhancing insulin (5; 6). As a key transcriptional target of PPAR_γ_, Adpn moreover accounts for many of the primary metabolic effects of PPAR_γ_-activating agents such as thiazolidinediones (TZDs) (7; 8) and FGF21 (9). However, clinical use of recombinant Adpn has been limited by the difficulties encountered in producing stable HMW oligomers, which require complex post-translational modifications that are technically challenging to reproduce (10).

EVs are lipid bilayer-enclosed structures that transport bioactive molecules, including proteins, lipids, and nucleic acids (11; 12). Their high bioavailability, biocompatibility, and low immunogenicity make them promising therapeutic delivery nanocarriers (12; 13). Beyond their role as passive carriers, EVs offer a unique opportunity to present membrane-associated ligands in a spatially organized and biologically relevant configuration, a feature that is difficult to achieve with recombinant proteins (12; 14).

Adipocyte-derived extracellular vesicles (EVs) naturally carry multimeric Adpn on their surface (15; 16). We previously showed that this EV-associated Adpn accounts for ~20% of circulating Adpn and exhibits higher stability in the bloodstream than free Adpn, while retaining the insulin-sensitizing properties of the adipokine through AdipoR signaling (15).

Here, we move beyond descriptive biology to develop a bioengineered EV-based strategy that enables to use EVs as delivery vehicles for bioactive Adpn oligomers, paving the way for their use as a therapeutic platform. To this goal, we engineered EVs presenting anchored oligomerized Adpn forms on their surface (EV^PP-Adpn^) to target metabolic tissues and evaluate their potential in mitigating insulin resistance and metabolic dysfunctions in high-fat diet (HFD)-fed mice. Six weeks of EV^PP-Adpn^ treatment improve insulin sensitivity, promote adipocyte lipid storage through insulin-regulated lipogenesis and alleviate key features of MASH lesions (steatosis, inflammation, fibrosis) by restoring hepatic sensitivity. Altogether, our results highlight that surface Adpn-exposing EVs as a promising biotherapy for managing T2D and MASH.

## Methods

For detailed methods, please refer to Supplementary material.

### Animal experimentation

Male and female C57BL/6J (B6) mice (10 weeks old) were fed a HFD (D12492, Safe Diets) *ad libitum* for 6 weeks and housed under standard conditions. Mice received biweekly intraperitoneal injections of EVs (EV^CTRL^ lacking Adpn or EV^PP-Adpn^) resuspended in 100 µL sterile 0.9% NaCl, or an equivalent volume of Vehicle (PBS in 0.9% NaCl). A 25 ng Adpn-equivalent EV dose, or the corresponding particle numbers for EV^CTL^, was selected based of our previous work, in which 5 µg injected adipose-derived EVs delivered ~5 ng Adpn and ~5×10^9^ particles per injection as measured by NTA (ZetaView) (15). In the present study, the 25 ng Adpn-equivalent dose corresponds to ~5-6× 10□ particles by NanoFCM and ~1 × 10□ by NTA, i.e. a particle number in the same order of magnitude but carrying a higher Adpn load (see Supplementary Fig.1I).

During the fifth week of EV injections, glucose (GTT) and insulin (ITT) tolerance tests were performed after a 4 hours of fasting. For GTT, mice received 1.25 g/kg glucose i.p., and for ITT, 0.75 IU/kg human insulin (Humulin). Blood glucose was measured from tail vein samples up to 120 min (Accu-Chek Guide).

At sacrifice, body and tissue weights were recorded in random-fed conditions or after a 4 h fast followed by insulin injection (3 IU/kg; Humulin, Lilly). Tissues were fixed for histology or snap-frozen in liquid nitrogen and stored at –80□°C for molecular analyses. All procedures were approved by the French Ministry of Research and the local ethics committee (CEEA - 006) and complied with EU Directive 2010/63/EU.

### Quantification and Statistical Analysis

Data were analyzed with GraphPad Prism and presented as dot plots of independent experiments. Statistical tests used are specified in figure legends. Significance was set at p ≤ 0.05, indicated as *p ≤ 0.05, **p ≤ 0.01, ***p ≤ 0.005, and ****p ≤ 0.001.

## Results

### Generation and characterization of bioengineered EVs carrying Adpn

To exploit the therapeutic potential of surface Adpn-associated EVs, we engineered EVs carrying native membrane-anchored Adpn using a patented technology (see ESM methods). A chimeric construct (PP-Adpn) fused Adpn to a transmembrane domain (TM) and a pilot peptide (PP) that directs it to the EV secretion pathway (Supplementary Fig.1A). EV^PP-Adpn^ were isolated from the supernatants of stably transfected PP-Adpn-HEK 293T cells, while control EVs (EV^CTL^) were obtained from non-transfected cells (see Supplementary material). Both EV preparations were enriched in EV markers Alix, syntenin-1 and CD81, exhibited similar size distributions with a mean diameter of ~70 nm and yielded EV concentrations of approximately 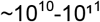 particles/mL (Supplementary Fig.1B-F). Adpn was consistently loaded in bioengineered EV^PP-Adpn^ batches, absent in EV^CTL^, and detected as high-molecular-weight, metabolically active forms (Supplementary Fig.1G-H).

### Surface Adpn-anchored EVs (EV^PP-Adpn^) improve insulin sensitivity

Male and female HFD-fed mice were injected twice weekly for 6 weeks with Vehicle (PBS), EV^CTL^ or EV^PP-^Adpn to assess their metabolic effects, following our EV-injection protocol that previously revealed the beneficial effects of vesicular Adpn from adipocyte-derived EVs (15). No significant differences in body weight gain over time (Fig.1A) or tissue weights (adipose depots, liver, pancreas, spleen and muscle; Supplementary Fig.2A) were observed. Blood biochemistry analyses (Supplementary Table 1) showed no hepatic or signs of renal toxicity, with even a slight trend toward lower transaminase levels in HFD-fed mice injected with EV^CTL^ compared to Vehicle group, supporting the absence of toxicity and confirming the good tolerance of repeated EV injections.

**Figure 1.**
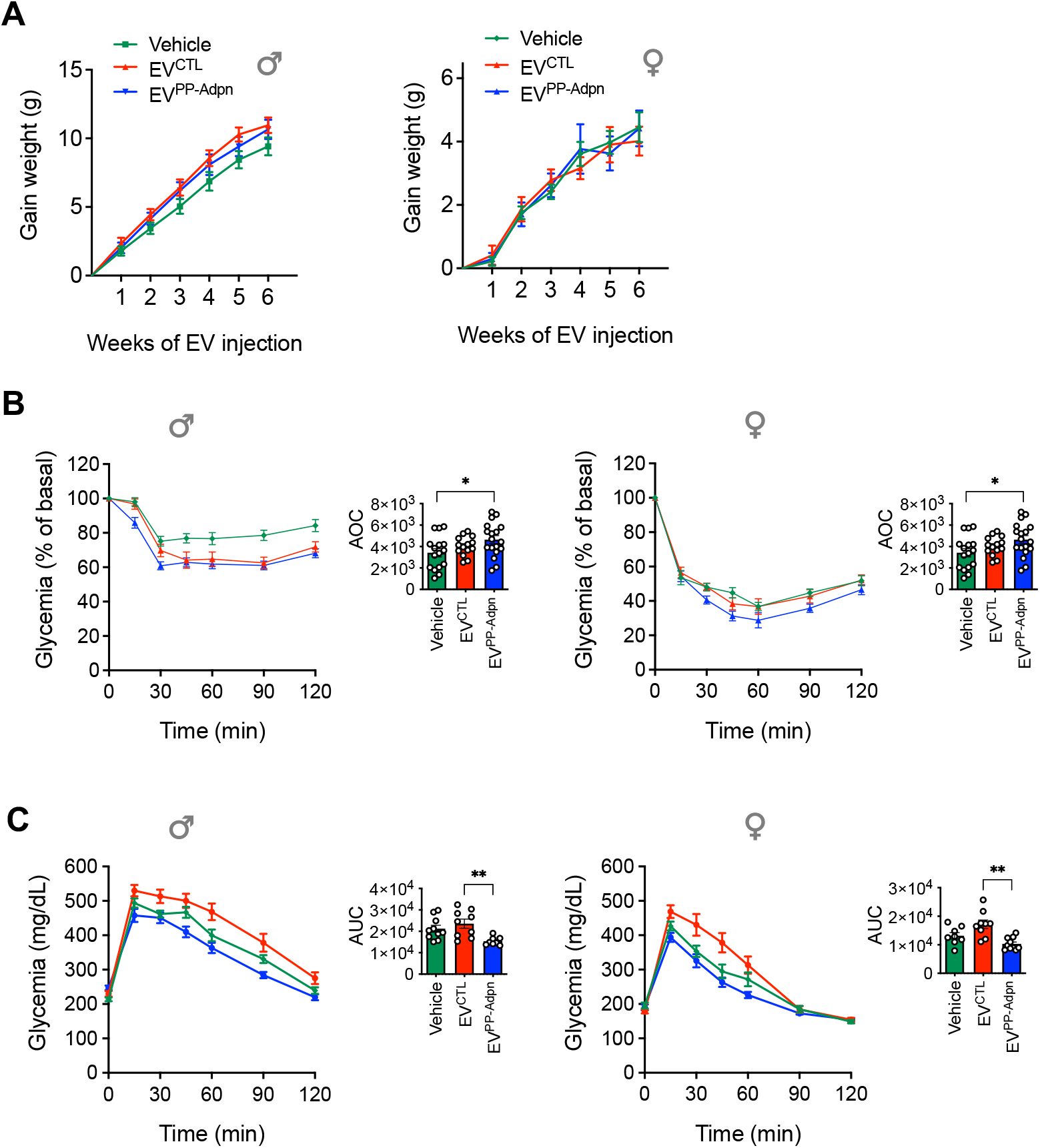
Surface-anchored Adpn-EVs (EV^PP-Adpn^) improve insulin sensitivity and glucose tolerance in HFD-fed mice. **(A)** Weight gain trajectory male (left) and female (right) HFD-fed mice after 6 weeks of i.p. injections with EV^CTL^ and EV^PP-Adpn^ or vehicle (PBS). **(B)** Insulin tolerance tests (ITT) performed in males (left) and females (right), with area over the curve (AOC) presented on the right. **(C)** Intraperitoneal glucose tolerance tests (GTT) in males (left) and females (right), with area under the curve (AUC) shown on the right.

Although random-fed and fasting glycemia remained stable in both sexes (Supplementary Fig.2B–C), EV^PP-Adpn^ significantly improved insulin sensitivity in males and females compared to Vehicle, as shown by increased area over the curve in ITT (Fig.1B). IP-GTT further demonstrated enhanced glucose tolerance in EV^PP-Adpn^-treated mice versus EV^CTL^ (Fig.1C). In males, this effect was accompanied by restoration of the 15-min insulin peak after glucose load, absent in EV^CTL^, Vehicle-treated groups, and all females (Supplementary Fig.2D). These benefits occurred without changes in pancreatic β-cell mass or size, as confirmed by comparable insulin immunostaining across groups (Supplementary Fig.2E–H).

### EV^PP-Adpn^ promote healthy adipose tissue remodeling

Although total AT mass was unchanged (Supplementary Fig.2A), visceral adipose tissue (VAT) histology showed larger adipocytes in EV^PP-Adpn^ groups (Fig.2A-C), an effect not seen in subcutaneous adipose tissue (SAT, Supplementary Fig.2I-J). EV^PP-Adpn^ increased key adipocyte markers (*PPAR*_γ_*2, PLIN1, AdipoQ*) and insulin-regulated lipogenic enzymes (*ELOVL6, FAS, ACC, ACLY*) without affecting inflammatory gene expression (Fig.2D-F). Circulating non-esterified fatty acids (NEFA) levels were unchanged (Supplementary Table 1), indicating that basal lipolysis was unaffected. Together, these results show that surface Adpn-associated EVs promote insulin-sensitive VAT expansion and healthy adipose tissue remodeling.

**Figure 2.**
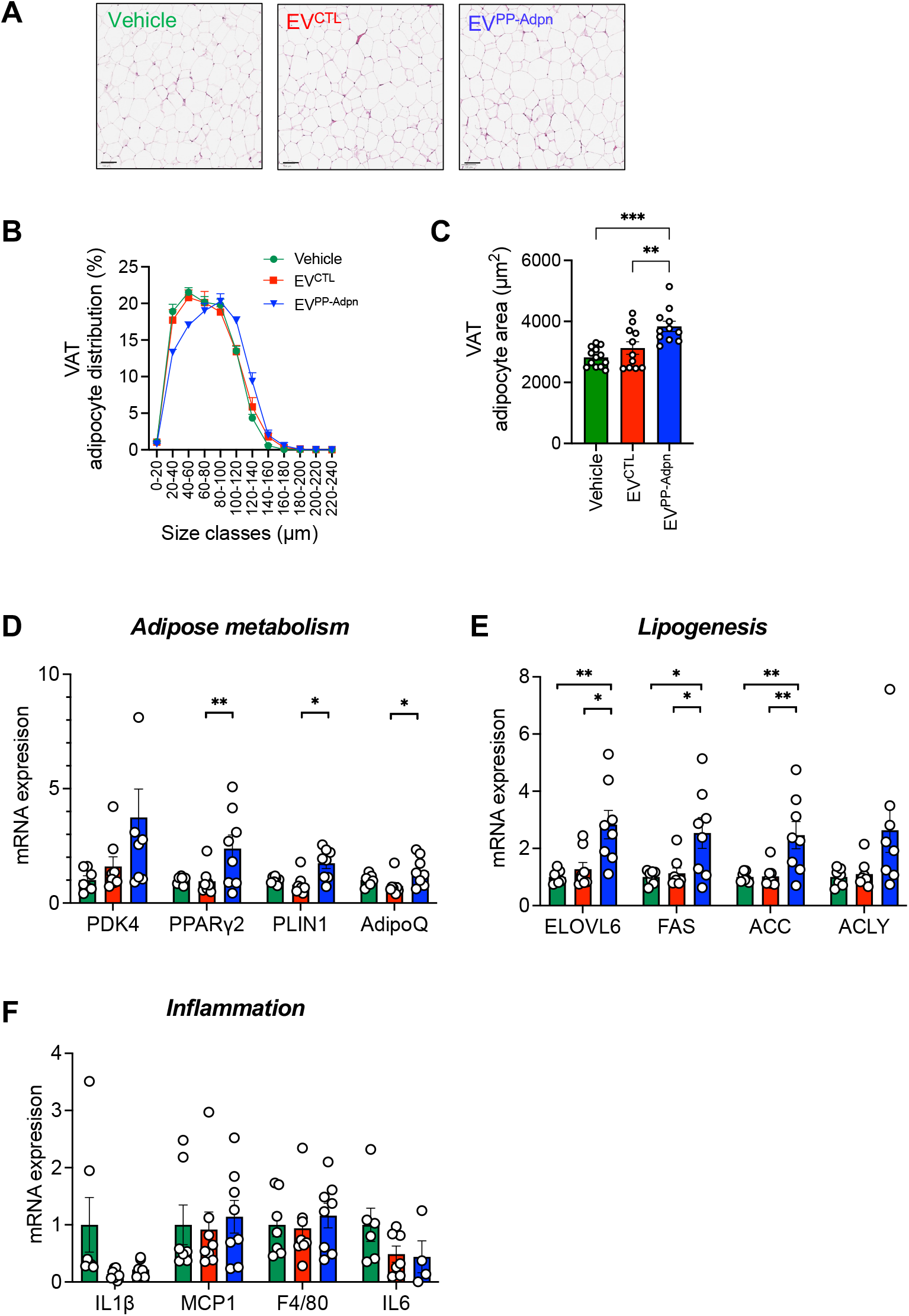
EV^PP-Adpn^ promote healthy adipose tissue remodeling. **(A-C)** Histological analysis of VAT in male HFD-fed mice at sacrifice. Representative hematoxylin/eosin-stained VAT sections are shown **(A)**, from which adipocyte diameters were measured. Analysis of VAT adipocyte size distribution is presented in **(B)**, and mean adipocyte area in **(C)**. n=11-12 males per group were analyzed. Scale bar, 100□µm. **(D-F)** Quantitive PCR gene expression analysis in VAT from male HFD-fed injected mice. mRNA expression was analyzed for the key adipogenic markers PDK4, PPAR_γ_2, PLIN1, AdipoQ **(D)**, for the insulin-responsive lipogenic genes ELOVL6, FAS, ACC, ACLY **(E)** and for the pro-inflammatory markers IL1β, MCP1, F4/80, IL6 **(F).** Data are presented as mean ± SEM. Dot plots represent the number of independent animals analyzed. Statistical differences were calculated using one-way ANOVA followed a Tukey’s multiple comparisons test. *p < 0.05, **p < 0.01. Bar colors: Green, Vehicle; Red, EV^CTL^; Blue, EV^PP-Adpn^.

### EV^PP-Adpn^ alleviate liver dysfunction and improve key metabolic signaling pathways

Liver histology revealed a marked reduction in steatosis in EV^PP-Adpn^–treated males, as highlighted by a significant decrease in hepatic triglyceride content compared to EV^CTL^ injected animals (Fig.3A; 3C-D). Liver fibrosis, already minimal in HFD-fed animals, remained unchanged (Fig.3B; 3E). Liver enzymes (ALT, AST) were significantly reduced in EV^PP-Adpn^ treated mice compared to Vehicle, whereas the modest decrease observed following EV^CTL^ remained non-significant (Supplementary Table 1). In contrast, EV^PP-Adpn^ markedly lowered circulating FGF21 compared with both EV^CTL^ and Vehicle, highlighting a specific Adpn-associated EV effect (Fig.3F). As circulating Fibroblast Growth Factor 21 (FGF21) and liver transaminases are typically elevated under HFD-induced metabolic stress, their coordinated reduction is consistent with improved hepatic function.

**Figure 3.**
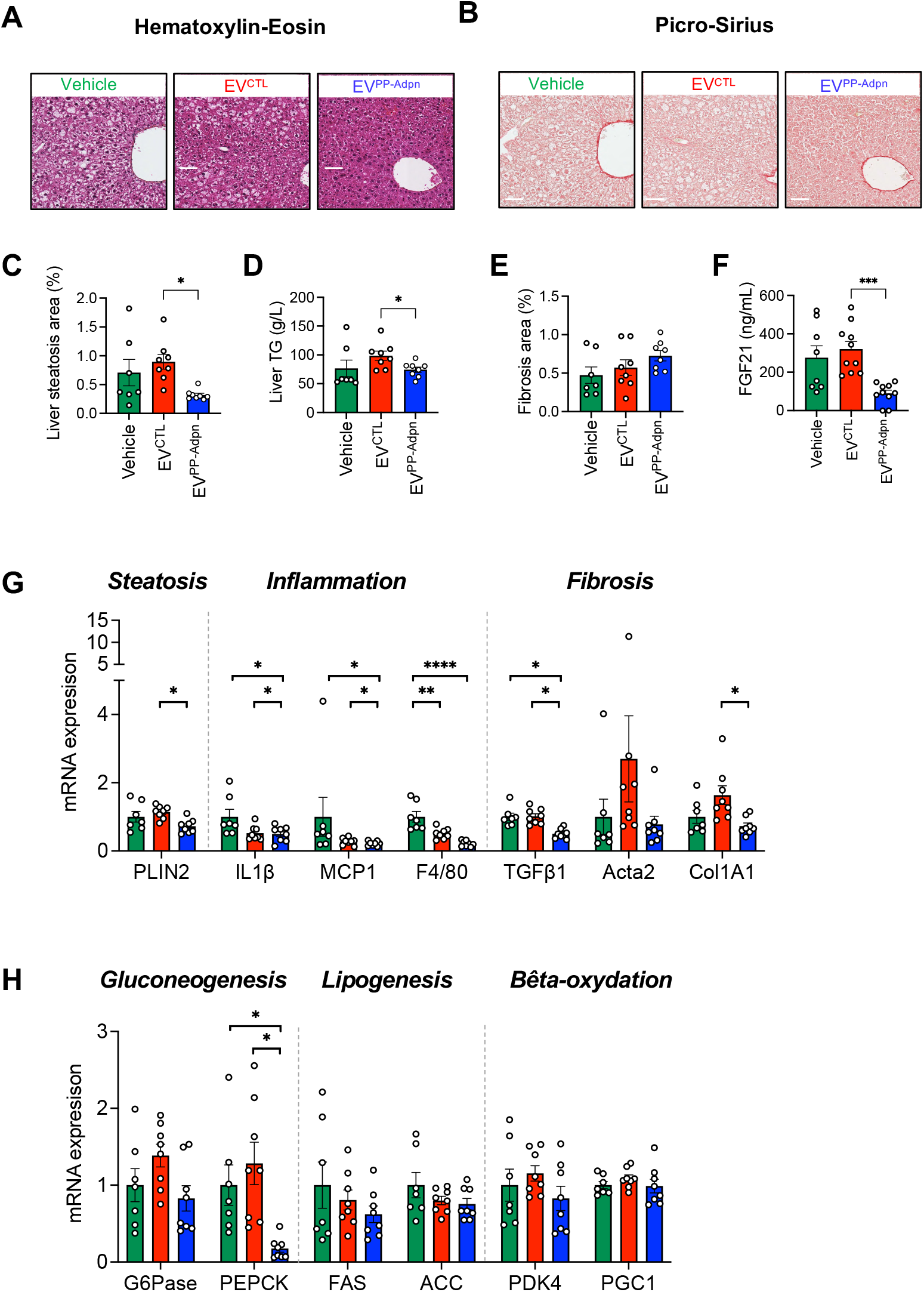
EV^PP-Adpn^ alleviate liver dysfunction and improve key metabolic signaling pathways. **(A-E)** Histological analysis of liver sections. Representative hematoxylin–eosin **(A)** and picrosirius red **(B)** staining are shown for male mice after 6 weeks of treatment with EV^CTL^ or EV^PP-Adpn^ or vehicle (PBS). Quantification of steatosis area on HE-stained sections **(C)**, hepatic triglyceride content **(D)** and fibrosis area on PS-stained sections **(E)** is presented. n=6-8 animals per group were analyzed. Scale bar, 100□µm. **(F)** Circulating FGF-21 in injected male mice. **(G-H)** Quantitative PCR analysis of hepatic gene expression in HFD-fed injected male mice. Relative mRNA levels of MASLD-associated genes **(G)** including steatotic (PLIN2), pro-inflammatory (IL1β, MCP1, F4/80) and fibrotic markers (TGFβ1, Col1A1). Expression of key metabolic genes **(H)** related to gluconeogenesis (PEPCK, G6Pase), lipogenesis (FAS, ACC) and β-oxidation (PDK4, PGC1α).

At the transcriptional level, EV^PP-Adpn^ downregulated metabolic dysfunction–associated steatotic liver disease (MASLD)-related genes including the lipid droplet marker *PLIN2*, pro-inflammatory (IL1β, MCP1, F4/80) and fibrotic markers (TGFβ1, Col1A1) (Fig.3G). Although most hepatic metabolic genes remained unchanged, EV^PP-Adpn^ restored insulin-mediated repression of PEPCK, with a similar trend for G6Pase (Fig.3H). Overall, these findings indicate that EV^PP-Adpn^ improve hepatic insulin sensitivity and alleviates key features of MASH.

### EV^PP-Adpn^ improve tissue-specific insulin signaling via activation of the adiponectin pathway

We next assessed adiponectin-responsive signaling in VAT, liver and skeletal muscle of male mice. Insulin-stimulated AKT (Ser473) phosphorylation -a readout of insulin sensitivity-was increased in all tissues of EV^PP-Adpn^ treated mice compared to EV^CTL^ mice, with significant effects in liver and muscle (Fig.4A). Given the central role of AMPK in adiponectin-driven regulation of lipid metabolism and energy homeostasis, we assessed AMPK (Thr172) phosphorylation. EV^PP-Adpn^ promoted a general increase across tissues, reaching significance in muscle, indicative of improved metabolic function (Fig.4B).

**Figure 4.**
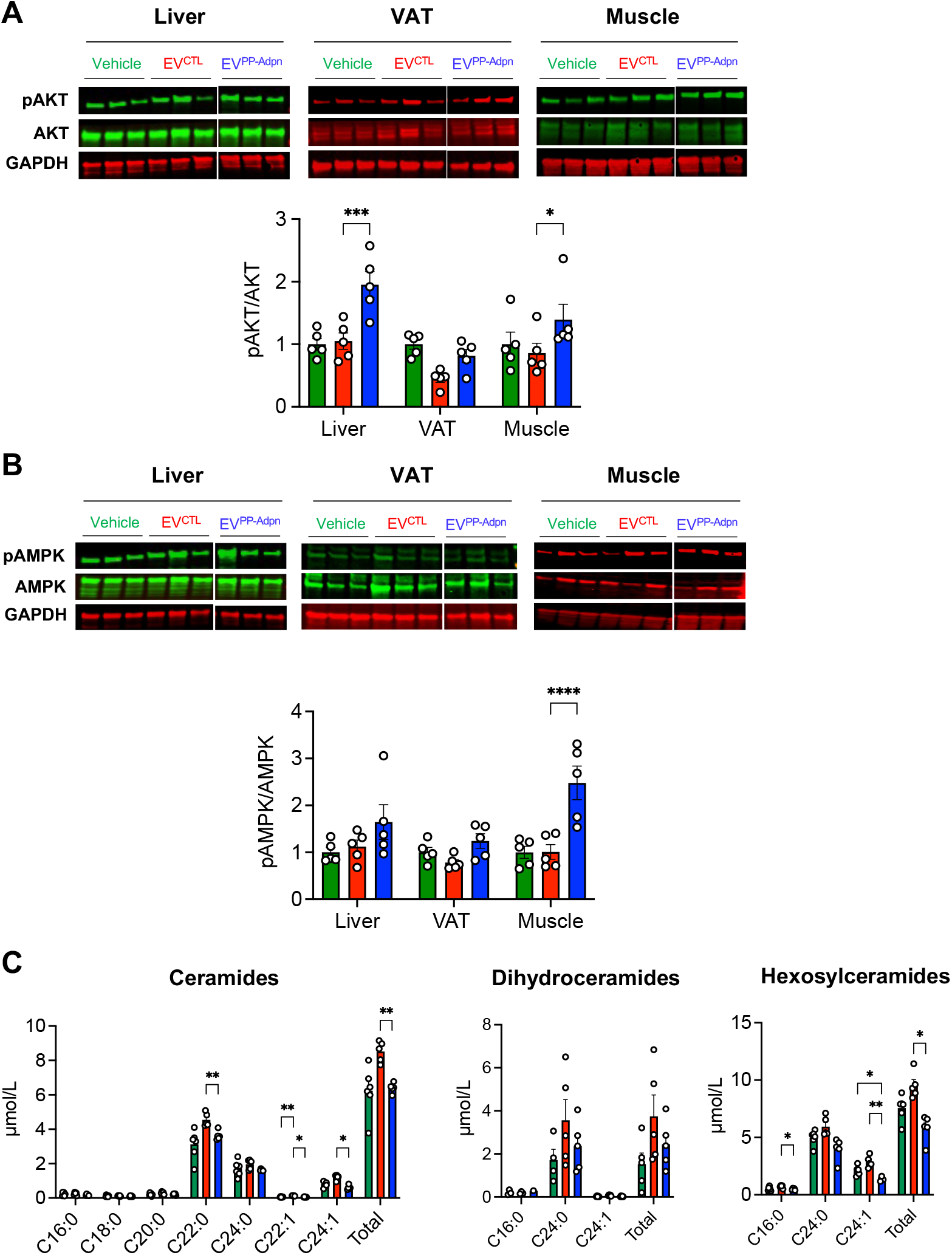
EV^PP-Adpn^ improve tissue-specific insulin signaling via activation of the adiponectin pathway. **(A-B)** Western blot analysis of AKT (Ser473) **(A)** and AMPK (Thr172) **(B)** phosphorylation in liver, VAT, and skeletal muscle following insulin injection in mice. Representative blots are shown, with GAPDH as loading control. Since EV^PP-Adpn^ and EV^CTL^ samples were run on the same gel but not in adjacent lanes, the non-adjacent lanes were cropped for clarity. Quantification of phospho-AKT/total AKT **(B)** and phospho-AMPK/total AMPK ratios **(B)** is presented below the blots. **(C)** Plasma ceramide profiling showing total levels and subspecies of ceramides (left panel), dihydroceramides (middle panel) and hexosylceramides (right panel). Data are presented as mean ± SEM. Dot plots represent the number of independent animals analyzed. Bar colors: Green, Vehicle; Red, EV^CTL^; Blue, EV^PP-Adpn^. icle

Because surface-anchored Adpn on EV^PP-Adpn^ is expected to activate AdipoR signaling (15), which promotes ceramidase activity, we measured plasma ceramides. EV^PP-Adpn^ treatment significantly decreased total ceramides and hexosyl-ceramides, particularly long-chain subspecies (C22:0 to C24:0) (Fig.4C). Altogether, these findings support that EV^PP-Adpn^ enhance tissue-specific insulin signaling through activation of adiponectin pathway, leading to metabolic improvements at both the systemic and organ levels.

## Discussion

EV^PP-Adpn^, engineered to display oligomeric adiponectin anchored at the vesicle surface, demonstrated insulin-sensitizing effects in HFD-fed mice, promoting adipocyte lipid storage and alleviating liver steatosis and inflammation. While the precise mechanism was not directly assessed in this study, anchoring of Adpn at the vesicle surface likely promotes AdipoR-mediated signaling, as suggested by our previous results with adipose-derived EVs (15). Although EVP^P-Adpn^ improve insulin sensitivity in both male and female mice, metabolic responses differ slightly as illustrated by the lack of an insulin secretion peak after a glucose challenge in females. These sex-specific differences may reflect distinct regulation of adiponectin signaling, as females have higher circulating adiponectin levels, potentially further enhanced by estrogen effects (17). The activation of the PPAR_γ_ pathway in VAT, the improvement in hepatic insulin sensitivity, and the activation of AMPK in skeletal muscle collectively demonstrate that the EV^PP-Adpn^ efficiently activate adiponectin signaling across these target tissues. As a consequence, the promotion of adipocyte lipid storage is consistent with enhanced insulin sensitivity and resembles the effects of other insulin-sensitizing agents, such as thiazolidinediones, whose metabolic actions have also been linked to adiponectin signaling (18).

Despite these benefits, EV^PP-Adpn^ were somewhat less potent than native adipose EVs, suggesting that additional vesicular cargos could act synergistically to further enhance insulin sensitivity (19). This concept is supported by several recent studies highlighting the therapeutic potential of engineered EVs with adipocyte-derived metabolic active molecules: leptin-loaded macrophage-derived EVs sucessfully targeted the brain and restored breathing function in obese mice, overcoming leptin resistance (20). Similarly to our bioengineered Adpn-enriched EVs, apelin-loaded EVs improved glucose tolerance and activated AMPK and Akt pathways in diabetic mice, enhancing systemic insulin sensitivity (21). Finally, EVs combining surface FGF21 with miR-223 achieved liver-preferential delivery and reduced the MASH phenotype (22). These examples highlight the potential of combinatorial cargo strategies to further potentiate EV^PP-Adpn^ and optimize therapeutic efficacy in insulin resistance and metabolic diseases.

## ACKNOWLEDGMENTS

We thank the SCAHU platform, especially E. Guillet, J. Roux, A. Mourlan, N. Meignan-Fizanne for mouse housing, care and technical assistance for animal functional exploration. We also thank the PACeM platform, especially L. Bonneau and J. Cayon, for qRT-PCR experiments. The graphical abstract was created with BioRender.com.

## AUTHORS’ CONTRIBUTIONS

Methodology for EV production and characterization, A.B., M.L., K.P., R.M., B.T.

Methodology for animal experimentation and analysis, J.F., M.V., G.H., L.M., J.M., L.F., X.P., C.L.M., B.C., S.L.L.

NTA analysis, J.F., M.V., S.L.L.

Histology analysis, Q.M., J.C.

Blood biochemistry analysis, M.C.

Lipidomic analysis, M.C., M.G.;

Conceptualization, S.L.L., B.C., R.M., B.T.

Writing – original draft, S.L.L.

Writing, review and editing, J.F., M.V., X.P., C.L.M., J.B., B.C., A.B., R.M., S.L.L.

Supervision of the animal experimental part and project coordination, S.L.L.

All authors have read and agreed to the published version of the manuscript.

S.L.L. is the guarantor of this work and, as such, had full access to all the data in the study and takes responsibility for the integrity of the data and the accuracy of the data analysis.

## Abbreviations

ACC: acetyl-CoA carboxylase
ACLY: ATP citrate lyase
AdipoQ: adiponectin gene
ALT: alanine aminotransferase
AST: aspartate aminotransferase
BAT: brown adipose tissue
Col1A1: collagen type I alpha 1
ELOVL6: elongation of very long chain fatty acids protein 6
F4/80: EGF-like module-containing mucin-like hormone receptor-like 1 (macrophage marker)
FAS: fatty acid synthase
FGF21: circulating Fibroblast Growth Factor 21
IL-1β: interleukin-1 beta
IL-6: interleukin-6
MCP1: monocyte chemoattractant protein 1
PDK4: pyruvate dehydrogenase kinase 4
PGC1α: peroxisome proliferator-activated receptor gamma coactivator 1-alpha
PLIN1: perilipin 1
PLIN2: perilipin 2
PPAR_γ_2: peroxisome proliferator-activated receptor gamma isoform 2
TGF-β1: transforming growth factor beta 1.

