## Supplemental Material for "Extracellular vesicles carrying surface-anchored adiponectin prevent obesity-related metabolic complications by enhancing insulin sensitivity"

**Supplementary Methods**

**Gene synthesis and molecular cloning**

Gene encoding adiponectin (accession number: NM_001177800.2), were codon optimized for human cells to generate Adiponectin gene. A PP-Adpn chimeric construct was generated encoding the adiponectin gene (Adpn) fused at its N-terminus to a sequence encoding transmembrane domain (TM) and to the sequence encoding a pilot peptide (PP, Patent WO2023104822A1, (1)). This construct was subcloned into a eukaryotic expression plasmid that also carries a zeocin resistance gene.

**Cell culture and EV production**

HEK293T were cultured in DMEM supplemented with 5% heat inactivated fetal bovine serum (iFBS), 2 mM GlutaMAX and 5 µg/mL of gentamicin at 37 °C in a 5% CO2 humidified incubator. HEK293T cells were transfected with adiponectin DNA using PEI. Zeocin (Invivogen) selection pressure was applied in order to establish stable cell lines. Stably transfected HEK293T cells expressing the PP-Adpn chimeric construct were cultured in cell chambers of 10 trays in complete medium and used for EV^PP-Adpn^ production. Twenty-four hours after, cultures were fed with an EV-free medium and incubated for a further 48 hours. Mock-transfected HEK293T cells were used for the production of control EVs (EV^CTL^) lacking adiponectin.

**EV purification**

Conditioned cell culture medium was harvested and the EV isolation was performed as previously described (2). Briefly, cell culture supernatant was clarified by two consecutive centrifugations: 10 minutes at 1300 rpm and 15 minutes at 4000 rpm, both at 4 °C, followed by filtration through 0.22 µm membrane filters. The supernatant was then concentrated by ultra-filtration/diafiltration and purified by Size Exclusion-based Chromatography. Fractions containing EV CD81 marker were identified by ELISA and pooled.

**EV size distribution and particle number**

EV size distribution and absolute size distribution were obtained by nanoflow cytometry (Nano-FC) using a NanoAnalyzer instrument (NanoFCM). EV concentrations were also measured by NTA using the ZetaView system to provide an additional quantification of particle numbers, as the two systems rely on different measurement principles. The results were validated through at least three replicates.

**SDS-PAGE, Western blotting and antibodies**

Protein concentrations of EV batches were determined using the BCA assay (Thermo Scientific). For the identification of EV markers (Alix and Syntenin-1) by SDS-PAGE, 5 μg of pure EVs were lysed in denaturing buffer and heated for 10 min at 95°C. For the detection of adiponectin, 1.25 μg of pure EVs were lysed in denaturing buffer and heated for 10 min at 95°C. Identification of adiponectin multimeric forms was performed in non-reducing and unheated conditions. EV preparations were separated by SDS-PAGE on a 4-15% gradient polyacrylamide gel (Bio-Rad) and proteins were subsequently transferred onto PVDF membrane. The immunodetection of proteins was performed with primary antibodies recognizing specifically either Alix (Proteintech), Adiponectin (Genetex) or Syntenin-1 (Fisher Scientific) proteins. Membranes were then incubated with the corresponding secondary Horseradish Peroxidase (HRP)-conjugated antibodies (donkey anti-goat HRP, donkey anti-mouse HRP or donkey anti-rabbit HRP, Jackson ImmunoResearch). The signals were detected using enhanced chemiluminescence detection.

Insulin and AMPK signaling were assessed in mouse tissues collected 15 minutes after i.p. insulin injection. VAT, SAT, and skeletal muscle (quadriceps) were harvested at this time point. Tissue homogenization was performed on 20mg of snap-frozen liver or muscle tissue and 100 mg of snap-frozen VAT using CK14 soft tissue beads and the Precellys homogenizer, according to the manufacturer’s instructions. Protein lysates (15 µg) were subjected to SDS–PAGE and Western blotting, as previously described¹¹. Primary antibodies used are phospho-Akt (Ser473, #4060), total Akt (pan, #4691), phospho-AMPKα (Thr172, #2535) and total AMPKα (#2532) (Cell Signaling Technology), GAPDH (#ABC-AC002-100, ABclonal), and β-actin (#A5316, Sigma-Aldrich). IRDye secondary antibodies and an Odyssey CLx imaging system (LI-COR) were use for detection and quantification was performed using Image Studio. Representative blots include samples from at least three independent animals per group.

**Anti-CD81 ELISA**

Serial dilutions of pure EVs (from 1 μg to 1 ng) were coated onto a 96-well ELISA plate overnight at 4°C. After saturation with 3% BSA in PBS during 1 h at 37°C, anti-CD81 was added and incubated for 2 h at 37°C. Then, the plate was washed three times and incubated with the corresponding secondary HRP-conjugated antibody for 1 h at 37°C. After washing 5 times, 3, 3', 5, 5' – Tetramethylbenzidine (TMB), the chromogenic peroxidase substrate, was added and the plate was incubated under dark for 30 min at room temperature. Sulfuric acid was added to stop the reaction. Optical density (OD) was measured at 450 nm using a CLARIOstar Plus plate reader (BMG Labtech).

**Quantitative anti-Adiponectin ELISA**

The adiponectin concentration was measured using commercial ELISA kits (Human Adiponectin DuoSet ELISA, R&D systems) designed to measure full-length human adiponectin (Adpn) levels according to the manufacturer’s protocol. 3 ng of Adpn EVs were lysed and diluted in 1X diluent reagent provided in the kit. The capture and HRP-conjugated detection antibodies as well as the calibrator provided in the total adiponectin ELISA kit were used. After substrate solution incubation, the optical density at 450 nm was read with a CLARIOstar Plus plate reader and corrected by the optical density read at 570 nm. Adpn concentrations were calculated from a four-parameter logistic (4-PL) standard curve fitting.

**Hepatic Lipid Quantification**

Hepatic lipids were extracted from snap-frozen liver samples using a modified Bligh and Dyer method (3). Triglycerides (TG) were quantified using enzymatic kits (Roche Diagnostics) and expressed relative to liver weight (µg TG per g of liver tissue).

**Histology and tissue staining**

Adipose and liver tissues were fixed in 10% PBS-buffered formalin for at least 24 hours, paraffin-embedded, and sectioned at 5 µm. After deparaffinization and rehydration with specific commercial solutions (Impath), sections were stained with with hematoxylin and eosin (HE) for morphology and picrosirius red (PS) to assess fibrosis. Whole-slide liver images were acquired using an Aperio digital slide scanner (Scanscope CS2 System, Aperio Technologies, Vista, CA, USA), providing high-resolution images (maximum scanning area capacity of 120,000 × 50,000 pixels at 0.5 μm/pixel, magnification ×20). Adipocyte diameter (VAT, SAT), steatosis and fibrosis areas (liver) were quantified using dedicated ImageJ plugins.

**RNA Extraction and qPCR**

RNA was extracted from liquid nitrogen–frozen tissues. For VAT, RNA was isolated using QIAzol and purified with the RNeasy Microkit (Qiagen), including DNase I treatment. Liver and muscle RNA were extracted using the Maxwell® RSC simplyRNA Tissue Kit and instrument (Promega). RNA quantity and purity were assessed via NanoDrop ND-2000 (Thermo Fisher). cDNA was synthesized from 1 µg RNA using SuperScript™ II (Invitrogen) and random hexamers, then purified with the Qiaquick PCR kit (Qiagen). qPCR was performed with 3 ng cDNA, Maxima™ SYBR Green Master Mix (Thermo Fisher), and 0.3 µM primers using a CFX Opus 384 system (Bio-Rad). Cycling conditions were 95°C for 10 min, then 40 cycles of 95°C for 15 s and 60°C for 30 s. Specificity was confirmed by melting curve analysis. Gene expression was calculated via the 2^−ΔΔCq^ method, normalized to 18S and 36B4, and analyzed with CFX Maestro (Bio-Rad). Primer sequences are available upon request.

**Ceramide analysis by mass spectrometry**

Ceramides were quantified from 10 µL of plasma using targeted lipid liquid chromatography–tandem mass spectrometry (LC-MS/MS), as previously described (4).

**Biochemical analysis of circulating metabolic markers**

Serum metabolic and hepatic parameters were measured from 100 µL of randomly fed mouse serum using a Cobas Pro analyzer (Roche Diagnostics). The following analytes were quantified using dedicated reagents: total protein, albumin, glucose, triglycerides, total cholesterol, HDL cholesterol, LDL cholesterol, alanine aminotransferase (ALT), aspartate aminotransferase (AST), and alkaline phosphatase (ALP). All measurements were performed according to the manufacturer’s protocols, with automatic internal quality control checks ensuring analytical reliability and reproducibility.

Circulating free fatty acids were measured using Non-Esterified Fatty Acid (NEFA) Assay Kit (Fujifilm) following the the manufacturer’s protocol.

Circulating FGF21 concentrations was measured using commercial ELISA kits designed for mouse measurments (R&D systems) from plasma samples according to the manufacturer’s protocol.

Plasma insulin during GTT was quantified by ELISA (Crystal Chem) from heparinized blood

**Supplementary Table 1: Blood biochemical profiles in male and female mice treated with bioengineered EVs.**

Data are presented as mean ± SEM for male (♂) and female (♀) mice treated with bioengineered EVs (25 ng Adpn-equivalent). The number of animals analyzed per group is indicated (*n*). Statistical differences were determined using one-way ANOVA followed by **Tukey’s multiple comparisons test.** * indicates p < 0.05 versus Vehicle, while ^###^ indicates p < 0.005 versus EV^CTL^. *ND*, not determined.

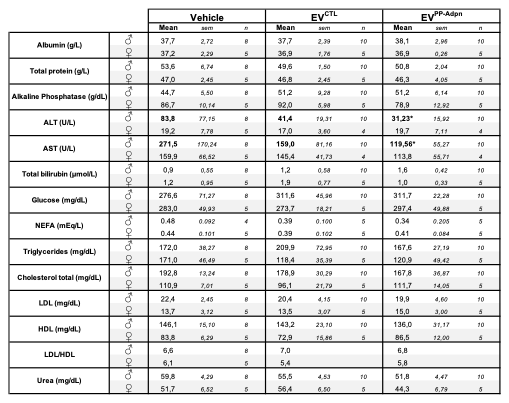

**Supplementary Fig.1. Design and validation of bioengineered EV-anchored Adpn.**

**(A)** Schematic representation of Adpn molecular construct used to establish a stable cell line expressing EV-anchored Adpn (EV^PP-Adpn^) via the fusion of adiponectin gene at its N-terminus to a sequence encoding a transmembrane domain (TM) and a pilot peptide (PP). Control EVs (EV^CTL^) lacking Adpn were produced from mock-transfected HEK293T cells.

**(B-C)** Enrichment of EV markers in bioengineered EVs. The presence of the EV marker proteins Alix and Syntenin-1 was assessed by Western blot **(B)**. Presence of the EV marker CD81 in EVs assessed by ELISA **(C)**. Standard EVs were used as positive EV marker controls (C+). Representative blots for both EV^CTL^ and EV^PP-Adpn^ are shown.

**(D-F)** Size distribution curves **(D)**, EV mean size **(E),** and EV concentration **(F)** of bioengineered EVs, as measured using a NanoFCM instrument (Nano-FC). EV concentrations were additionally assessed by NTA (ZetaView) in panel F. Dot plots represent independent production batches for each EV type. In Panel F, results are presented as mean ± sem (n=4 for EV^CTL^, n=3 for EV^PP-Adpn^).

**(G)** Adpn content was quantified by ELISA and expressed as ng Adpn per µg EV protein. Dot plots represent independent production lots per EV type analyzed.

**(H)** Western-blot analysis of EV-anchored Adpn (EV^PP-Adpn^) under reducing and non-reducing **(**conditions, demonstrating the presence of Adpn and its assembly under high-molecular-weight proteo-oligomeric forms.

**(I)** Corresponding injected EV numbers for EV^PP-Adpn^ and EV^CTRL^ at the 25-ng Adpn-equivalent dose, as quantified by Nano-Flow cytometry (NanoFC) or by NTA (ZetaView). Results are presented as mean ± sem (n=3 for EV^CTL^, n=2 for EV^PP-Adpn^).

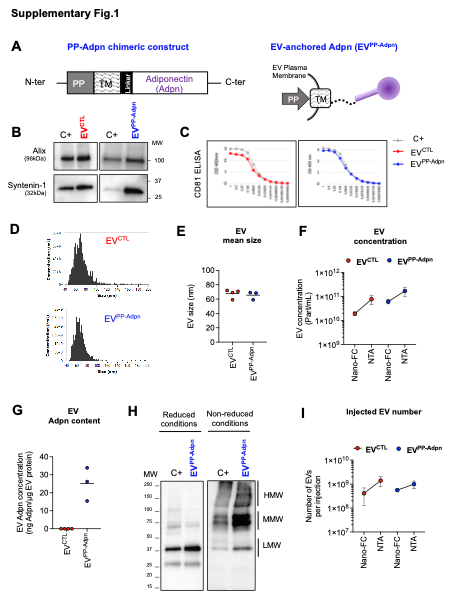

**Supplementary Fig.2. Metabolic phenotyping of EV^CTL^, EV^PP-Adpn^ or Vehicle injected HFD-fed mice**

**(A)** Tissue weight (in g) at sacrifice is presented for liver, VAT, SAT, BAT, pancreas, spleen and muscle following EV injections in males (left panel) or in females (right panel). Floating bars (min to max) with line at mean are presented for each condition.

**(B-C)** Random-fed glycemia measured throughout the experiment, n=16-18 for males (top); n=9-10 for females (bottom) **(B)** and fasting glycemia measured at sacrifice **(C)** in males (top) and females (bottom).

**(D)** Insulin levels measured at baseline (T0) and 15 minutes post-glucose injection (T15) during GTT in males (top) and females (bottom). Five animals per group were analyzed.

**(E-H)** Representative pancreatic sections stained for insulin showing comparable β-cell mass across treatment groups in male mice **(E)**. Quantification of β-cell fraction (%) **(F)**, mean islet size (µm²) **(G)**, and islet density (n/mm²) **(H)** are shown. Scale bar, 2 nm.

**(I-J)** Histological analysis of SAT sections in male (upper panels) and female (lower panels) mice following the EV injection protocol showed no significant changes in adipocyte size distribution **(I)** or mean adipocyte area **(J)** across treatment groups.

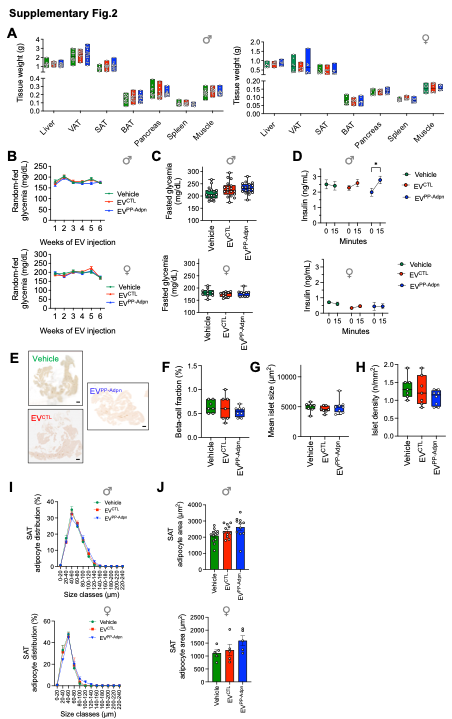
